# Accurate prediction of human essential genes using only nucleotide composition and association information

**DOI:** 10.1101/084129

**Authors:** Feng-Biao Biao, Chuan Dong, Hong-Li Hua, Shuo Liu, Hao Luo, Hong-Wan Zhang, Yan-Ting Jin, Kai-Yue Zhang

## Abstract

Three groups recently identified essential genes in human cancer cell lines using wet experiments, and these genes are of high values. Herein, we improved the widely used Z curve method by creating a λ-interval Z curve, which considered interval association information. With this method and recursive feature elimination technology, a computational model was developed to predict human gene essentiality. The 5-fold cross-validation test based on our benchmark dataset obtained an area under the receiver operating characteristic curve (AUC) of 0.8814. For the rigorous jackknife test, the AUC score was 0.8854. These results demonstrated that the essentiality of human genes could be reliably reflected by only sequence information. However, previous classifiers in three eukaryotes can gave satisfactory prediction only combining sequence with other features. It is also demonstrated that although the information contributed by interval association is less than adjacent nucleotides, this information can still play an independent role. Integrating the interval information into adjacent ones can significantly improve our classifier’s prediction capacity. We re-predicted the benchmark negative dataset by Pheg server (https://cefg.uestc.edu.cn/Pheg), and 118 genes were additionally predicted as essential. Among them, 21 were found to be homologues in mouse essential genes, indicating that at least a part of the 118 genes were indeed essential, however previous experiments overlooked them. As the first available server, Pheg could predict essentiality for anonymous gene sequences of human. It is also hoped the λ-interval Z curve method could be effectively extended to classification issues of other DNA elements.

## Introduction

Catalogs of essential genes on a whole-genome scale, determined using wet-lab methods, are available for several prokaryotic (Mushegian and Koonin 1996; Luo et al. 2014; Pechter et al. 2016) and eukaryotic organisms (Kamath et al. 2003; Amsterdam et al. 2004; Liao and Zhang 2007; Kim et al. 2010). Computational methods with high accuracy offer an appealing alternative method for identifying essential genes. Computational methods are broadly divided into three types: machine learning-based methods combining intrinsic and context-dependent features (Deng et al. 2011; Cheng et al. 2013), flux balance analysis-based methods (Kuepfer et al. 2005; del Rio et al. 2009; Gatto et al. 2015), and homology search and evolutionary analysis-based methods (Peng et al. 2012; Wei et al. 2013). With respect to essential gene prediction in bacteria, we integrated the orthology and phylogenetic information and subsequently developed a universal tool named Geptop (Wei et al. 2013), which has shown the highest accuracy among all state-of-the-art algorithms.

Some studies have focused on essential gene prediction in eukaryotic genomes. In 2005, Xu et al. investigated protein dispensability in *Saccharomyces cerevisiae* by combining high-throughput data and machine learning-based methods (Chen and Xu 2005). In 2006, Seringhaus et al. reported a machine learning-based method that integrated various intrinsic and predicted features to identify essential genes in yeast *S. cerevisiae* genomes (Seringhaus et al. 2006). They for the first time using AUC to measure the machine learning classifier’s capability in identifying essential genes, and they also translated the essentiality from yeast *S. cerevisiae* to yeast *Saccharomyces mikatae.* Yuan et al. integrated informative genomic features to perform knockout lethality predictions in mice using three machine learning-based methods (Yuan et al. 2012). Lloyd et al. analyzed the characteristics of essential genes in the *Arabidopsis thaliana* genome and used *A. thaliana* as a machine learning-based model to transform the essentiality annotations to *Oryza sativa* and *S. cerevisiae* (Lloyd et al. 2015).

Recently, three research teams approximately identified 2,000 essential genes in human cancer cell lines using CRISPR-Cas9 and gene-trap technology (Blomen et al. 2015; Hart et al. 2015; Wang et al. 2015). Their results showed high consistency, which further confirmed the accuracy and robustness of the essential gene sets (Fraser 2015). These studies provided an in-depth analysis of tumor-specific essential genes and feasible methods to screen tumor-specific essential genes (Fraser 2015; Hart and Moffat 2016). The essential genes screened by these three teams provided a clear definition of the requirements for sustaining the basic cell activities of individual human tumor cell types. Practically, these genes can be regarded as targets for cancer treatment (Fraser 2015). The data from these three groups provided a rare opportunity to theoretically study the function, sequence composition, evolution and network topology of human essential genes. One of the most important and interesting theoretical issues in modern biology is whether essential genes and non-essential genes can be accurately classified using computational methods. The models established in the aforementioned three eukaryotic organisms, *S. cerevisiae* (Chen and Xu 2005; Seringhaus et al. 2006), *Mus musculus* (Yuan et al. 2012), and *A. thaliana* (Lloyd et al. 2015), involved intrinsic features, or intrinsic and context-dependent features. These context-dependent features included those features extracted from experimental omics data. However, the features derived from experimental data are frequently unavailable; consequently, this type of machine learning model cannot be extended to a wide range of genomes. In the present study, we addressed this problem in humans by using only intrinsic features derived from sequences, from which certain features can be characterized using a λ-interval Z-curve. To facilitate the use of these data by interested researchers, we have provided a user-friendly online web server, Pheg, which can be freely accessed without registration at http://cefg.uestc.edu.cn/Pheg.

## Results

### Cross-validation Results

The final features of this method were described by *FV*_*w,λ*_, a value that contains information on the composition of the adjacent w-nucleotides (Gao and Zhang 2004) and λ-interval nucleotides. The association information was also captured by *FV*_*w,λ*_. Therefore, this method achieves improved performance compared with using the original Z-curve. The following results solidly confirmed this point.

We performed a 5-fold cross-validation test with w ranging from 2 to 4 and λ ranging from 0 to 5. The detailed results are provided in Table 1, showing that area under the curve (AUC) values gradually increased with increasing λ when w was fixed. An examination of the performance under variable values for w and fixed values for λ revealed that the performance for the classifier improved with increasing values for w.

**Table 1.** AUC values at different w, λ and penalty parameters c.

| Variables | $c$ | AUC |
| --- | --- | --- |
| $FV_{2,0}$ | 256 | 0.8002 |
| $FV_{2,1}$ | 64 | 0.8198 |
| $FV_{2,2}$ | 16 | 0.8256 |
| $FV_{2,3}$ | 128 | 0.8275 |
| $FV_{2,4}$ | 64 | 0.8293 |
| $FV_{2,5}$ | 64 | 0.8365 |
| $FV_{3,0}$ | 32 | 0.8276 |
| $FV_{3,1}$ | 1 | 0.8333 |
| $FV_{3,2}$ | 0.5 | 0.8347 |
| $FV_{3,3}$ | 0.25 | 0.8356 |
| $FV_{3,4}$ | 0.5 | 0.8408 |
| $FV_{3,5}$ | 0.25 | 0.8429 |
| $FV_{4,0}$ | 1 | 0.8344 |
| $FV_{4,1}$ | 0.5 | 0.8369 |
| $FV_{4,2}$ | 0.5 | 0.8386 |
| $FV_{4,3}$ | 0.25 | 0.8413 |
| $FV_{4,4}$ | 0.25 | 0.8436 |
| $FV_{4,5}$ | 0.25 | 0.8449 |

As shown in Table 1, we obtained an AUC value of 0.8002 under *FV*_*2,0*_. However, after utilizing the λ-interval nucleotide composition, the performance was improved, for example, the best AUC achieved for this model was 0.8449 through the 5-fold cross-validation test under variable *FV*_*4,5*_. The AUC was improved 4.47% compared with *FV*_*2,0*_. The information redundancy and noise in the original features can influence the predictive power of a classifier, and high-dimensional features also increase the time costs for training and prediction. Feature selection technology can mitigate these disadvantages. The support vector machine (SVM)-recursive feature extraction (RFE)+correlation bias reduction (CBR) method was adopted to rank these features in descending order based on the contribution of each feature. Subsequently,the top 100 features were used to constitute the initial feature subset to train and test the model, and the next 100 features were added into the feature subset, followed by prediction using the same methods. This process was repeated until the top 4,500 features had been added according to the rank order. The test results of each model were evaluated according to the AUC scores via a 5-fold cross-validation test. The AUC values for different top features are shown in Figure 1 (A).

**Figure 1.**
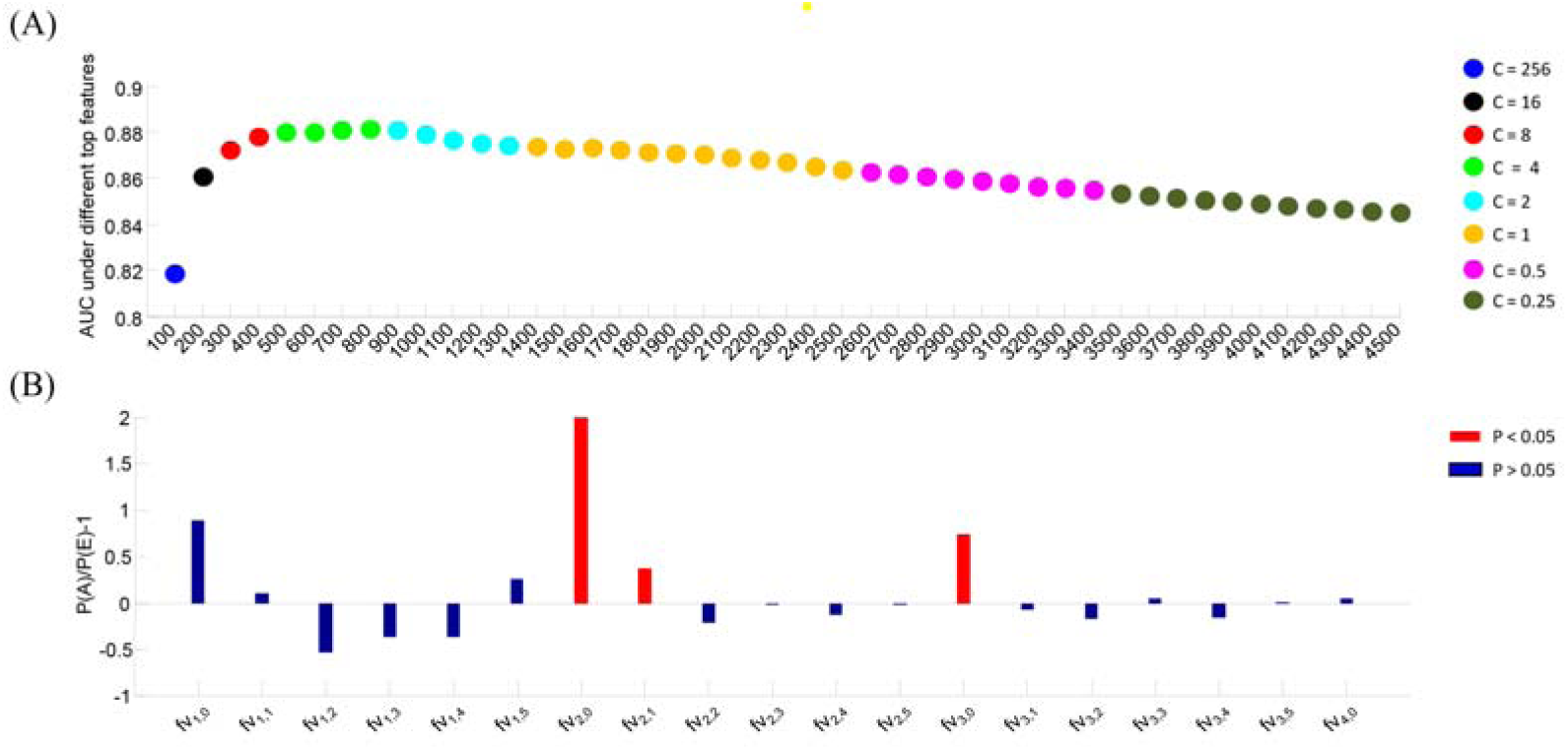
AUC values obtained under different top n features and the contribution of each group. Figure 1 (A) represents the AUC scores under different top features. Dots with different colors denote different c values. Figure 1 (B) illustrates the selective tendentiousness for every variable type. The red bars denote that the selective tendentiousness has significance.

Among all 4,545 features examined, the best AUC of 0.8814 was achieved for the top 800 selective features. The final AUC value was 8.12% higher than that for *FV*_*2,0*_. To conduct an objective evaluation of this method, we performed a rigorous jackknife test based on the top 800 selected features using the parameters determined via a 5-fold cross-validation test. We obtained an AUC value of 0.8854. As expected, excellent performance was obtained after adopting the λ-interval nucleotide composition and feature selection technology. Those results illustrated that the essentiality of human genes could be well reflected by only sequence information.

As we are extremely interested in the actual essential genes in the predicted results, we used the positive predictive value (PPV) to further refine. This evaluation index can be calculated using the formula TP / (TP+FP), where TP (true positive) and FP (false positive) represent the number of real essential and non-essential genes among the positive predictions. Therefore, the PPV reflects the proportion of actual essential genes among the predicted essential genes. We obtained a PPV of 73.05% (TP=515, FP=190) using the jackknife test based on the top 800 features. One of the simplest cross validation tests is the holdout method. In this procedure, the dataset is separated into two subsets, namely, training and testing datasets. We randomly sampled one-fifth of the positive and negative samples from the benchmark dataset for the training model, and the remaining samples were used as the testing dataset. To comprehensively assess the method used in the present study, we repeated the holdout method 100 times to identify differences in the composition of the training and testing samples. The mean AUC score was used as the final evaluator. A mean AUC score of 0.8537 with a variance of 1.67e-005 was obtained. Additionally, the proportions of samples in the training and predicting datasets were changed for further investigation. One-tenth of the positive and negative samples were randomly sampled as the training dataset, and the remaining samples were used as the testing dataset. This procedure was repeated 100 times. We obtained a mean AUC score of 0.8347 with a variance of 2.77e-005. These results further confirmed that this method was robust and accurate.

### Different contributions of each group of variables to the classification ability

Features with fixed w and λ values correspond to a specific group of variables. A total of 19 special groups were obtained, namely, *fv*_*1,0*_, *fv*_*1,1*_, *fv*_*1,2*_ … *fv*_*1,5*_; *fv*_*2,0*_, *fv*_*2,1*_ … *fv*_*2,5*_; *fv*_*3,0*_, *fv*_*3,1*_ … *fv*_*3,5*_, and *fv*_*4,0*_. We calculated the percentage of features in these groups, and the results are provided in Table 2.

For each group, there were two frequencies: P(A), which denotes the actual frequency of features in each group appearing in the top 800 selected features, and P(E), which denotes the expected frequency of the features in each group appearing in the original 4,545 features. Therefore, P(A) was obtained based on the number of selected features in each group divided by 800, and P(E) was calculated by dividing the number of total features in each group by 4,545. P(A) and P(E) are listed in columns 4 and 5, respectively. If P(A) is higher than P(E), then the group makes a higher-than-average contribution to the identification of essential genes. We calculated the tendentiousness using the formula P(A)/P(E)−1 to denote how the features of each group were selected, and the results are listed in column 6 of Table 2. We further conducted a hypergeometric distribution test for each group, and the p values are listed in column 7. Figure 1 (B) and Table 2 show that *fv*_*2,0*_ (p = 1.60E−06), *fv*_*2,1*_ (p = 0.02407688), and *fv*_*3,0*_ (p = 7.89E−05) are preferentially selected and are statistically significant. These results demonstrated that there are strong signals for classifying essential and non-essential genes when the character interval is equal to zero or one, but the other groups did not show these strong signals. To further confirm this result, the variables *fv*_*2,0*_, *fv*_*2,1*_, *fv*_*2,2*_, *fv*_*2,3*_, *fv*_*2,4*_, *fv*_*2,5*_ and *fv*_*3,0*_, *fv*_*3,1*_, *fv*_*3,2*_, *fv*_*3,3*_, *fv*_*3,4*_, and *fv*_*3,5*_ were used as input features. Improved performance was obtained under *fv*_*2,0*_, *fv*_*2,1*_, and *fv*_*3,0*_ compared with the other groups (Table 1, column 8). Those results demonstrated that the shorter interval association provides more information. However, longer interval association can still play an independent role. Hence, integrating the interval information into adjacent ones could significantly improve our classifier’s capacity of discernment (Table 1).

**Table 2.** Feature details for every variable type in the top 800 selective features. The feature groups with significance are indicated bold font.

| Variables | No.(A) | No.(E) | P(A) | P(E) | P(A)/P(E)-1 | p value | AUC | c |
| --- | --- | --- | --- | --- | --- | --- | --- | --- |
| $fv_{1,0}$ | 3 | 9 | 0.0038 | 0.0020 | 0.8938 | 0.200495 | — | — |
| $fv_{1,1}$ | 7 | 36 | 0.0088 | 0.0079 | 0.1047 | 0.452683 | 0.7220 | 1024 |
| $fv_{1,2}$ | 3 | 36 | 0.0038 | 0.0079 | -0.5266 | 0.965324 | 0.6216 | 64 |
| $fv_{1,3}$ | 4 | 36 | 0.0050 | 0.0079 | -0.3688 | 0.900314 | 0.6028 | 1024 |
| $fv_{1,4}$ | 4 | 36 | 0.0050 | 0.0079 | -0.3688 | 0.900314 | 0.6018 | 128 |
| $fv_{1,5}$ | 8 | 36 | 0.0100 | 0.0079 | 0.2625 | 0.292801 | 0.6302 | 256 |
| **$fv_{2,0}$** | **19** | **36** | **0.0238** | **0.0079** | **1.9984** | **1.60E-06** | **0.7902** | **64** |
| **$fv_{2,1}$** | **35** | **144** | **0.0438** | **0.0317** | **0.3809** | **0.024077** | **0.7551** | **1024** |
| $fv_{2,2}$ | 20 | 144 | 0.0250 | 0.0317 | -0.2109 | 0.906339 | 0.6841 | 1024 |
| $fv_{2,3}$ | 25 | 144 | 0.0313 | 0.0317 | -0.0137 | 0.566007 | 0.6633 | 512 |
| $fv_{2,4}$ | 22 | 144 | 0.0275 | 0.0317 | -0.1320 | 0.802176 | 0.6856 | 1024 |
| $fv_{2,5}$ | 25 | 144 | 0.0313 | 0.0317 | -0.0137 | 0.566007 | 0.6816 | 1024 |
| **$fv_{3,0}$** | **44** | **144** | **0.0550** | **0.0317** | **0.7359** | **7.89E-05** | **0.8236** | **2** |
| $fv_{3,1}$ | 94 | 576 | 0.1175 | 0.1267 | -0.0729 | 0.821727 | 0.7652 | 8 |
| $fv_{3,2}$ | 84 | 576 | 0.1050 | 0.1267 | -0.1715 | 0.983389 | 0.7125 | 32 |
| $fv_{3,3}$ | 107 | 576 | 0.1338 | 0.1267 | 0.0554 | 0.272681 | 0.7168 | 1024 |
| $fv_{3,4}$ | 86 | 576 | 0.1075 | 0.1267 | -0.1518 | 0.970308 | 0.7027 | 64 |
| $fv_{3,5}$ | 103 | 576 | 0.1288 | 0.1267 | 0.0159 | 0.44446 | 0.6983 | 1024 |
| $fv_{4,0}$ | 107 | 576 | 0.1338 | 0.1267 | 0.0554 | 0.272681 | — | — |

### A web server for predicting essential genes in human cancer cell lines

To facilitate the use of these data by interested researchers, we constructed a user-friendly online web server named Pheg (<u>**P**</u>redictor of <u>**h**</u>uman <u>**e**</u>ssential <u>**g**</u>enes), which is freely accessible at http://cefg.uestc.edu.cn/Pheg. The Pheg algorithm is based on the λ-interval Z-curve. Additional parameters are not necessary, making this algorithm convenient to use. Pheg can predict whether a query gene is essential using only the CDS region of a gene as input. We integrated logistic regression into the Pheg server to estimate the reliability of the predicted results. Hence, this server can output a probabilistic estimated value as a measurement of gene essentiality for the inputted coding region. Note that this is the first available server for predicting human gene essentiality. Comparatively, some computational models have been proposed for the other eukaryotes however all of them did not provide online prediction service. We re-predicted the genes in the benchmark dataset via Pheg and obtained an AUC = 0.9249 and PPV = 83.84%. A total of 612 genes were identified as essential genes among the 1,516 positive samples, and 118 genes were predicted as essential genes among the 10,499 negative samples. To estimate how many genes among those predictions are real essential genes, we calculated precisions using 5-fold, 10-fold, 15-fold, 20-fold cross-validation, and we obtained precisions with values of 70.43%, 71.63%, 72.48%, 72.22%, which were approximately 70%. Hence, we expect that 82 (118×70%) are correctly predicted essential genes. The information for these 118 genes is provided in Supplemental Information S1.

CancerResource is a comprehensive knowledgebase for drug-target relationships associated with cancer and for supporting information or experimental data (Gohlke et al. 2016). Through the CancerResource database (Gohlke et al. 2016), these 118 genes were searched using the classifier ‘Target search by gene identifiers’ (https://data-analysis.charite.de/care/index.php?site=target_input). Among these 118 genes, 17 genes showed interactions with drugs and other chemical molecules, and 13 genes had Cancer3D structures (Porta-Pardo et al. 2014). The details of these genes are supplemented in Supplemental Information S2. Additionally, we used these 118 gene sequences to conduct a BLAST (Basic Local Alignment Search Tool) search against essential genes in the genome of *Mus musculus* (mouse). The current mouse essential gene set is accessible in the OGEE database (Chen et al. 2012). Considering that no BLAST program is embedded in OGEE, we downloaded the essential gene annotations (gene_essentiality) at (http://ogee.medgenius.info/downloads/) and extracted the essential gene annotation of *M. musculus*. We obtained the essential gene sequences according to the annotations (http://ogee.medgenius.info/downloads/). A BLAST search was performed via ncbi-blast-2.2.30+-win64.exe using the data from OGEE, and homologs for 21 genes were identified (e value < 1e-100) among the 118 predicted essential genes. The sequences of the essential genes in *M. musculus* are provided in Supplemental Information S3. The details for these 21 genes and their corresponding homologs are provided in Supplemental Information S4. Both CancerResource analyses and BLAST search illustrated that at least a part of these 118 genes have higher probability to be factually essential genes and have been overlooked in the essential gene screening in previous experimental studies. Hence, Pheg sever could be used to predict essentiality for anonymous gene sequences of human and closely related species, and it is also hoped to supplement the essential gene list of human by identifying novel essential gene from the submitted sequences.

## Discussion

The Z-curve has been widely used in the field of bioinformatics for tasks such as protein coding gene identification (Zhang and Wang 2000; Chen et al. 2003; Guo et al. 2003; Guo and Zhang 2006; Hua et al. 2015), promoter recognition (Yang et al. 2008), translation start recognition (Ou et al. 2004), recombination spots recognition (Dong et al. 2016), and nucleosome position mapping (Wu et al. 2013). However, correlation and λ-interval nucleotide composition have not been incorporated into the Z-curve method. In the present study, we present a λ-interval Z-curve based on Z-curve theory. The DNA sequence can be understood as an ordinary character sequence; therefore, the method proposed in the present study has the potential for applications in mining characteristics from other character sequences and can be used as a universal feature extraction method for DNA sequences.

Based on the λ-interval Z-curve, we obtained excellent performance in human essential gene identification. This excellent performance might be attributable to the following points: First, we introduced the concept of intervals, reflecting associated information and the λ-interval nucleotide composition. Second, we used feature selection technology in the present study. Thus, noisy and redundant features could be removed from the original features. Table 2 shows the improved performance obtained under *fv*_*2,0*_, *fv*_*2,1*_, and *fv*_*3,0*_ compared with the other variable groups. Further comparison of these results with other feature groups shown in Table 1, and this comparison shows that the AUC values obtained with λ-interval variables are smaller than those obtained with shorter interval variables. However, the performance can be improved after adding λ-interval oligonucleotide association information (see Table 1). Hence, the λ-interval Z-curve should reflect additional important information for essential genes that cannot be contained in adjacent nucleotide association information.

In 2005, Chen and Xu used a neural network and SVM to predict the dispensability of proteins in the yeast *S. cerevisiae* based on the protein evolution rate, protein-interaction connectivity, gene-expression cooperativity and gene-duplication data (Chen and Xu 2005). The next year, Seringhaus et al. only used 14 features to predict essential genes in *S. cerevisiae* and obtained a PPV=0.69 (Seringhaus et al. 2006). Yuan et al. assembled a comprehensive list of 491 candidate genomic features to predict a lethal phenotype in a knockout mouse using three machine learning methods (Yuan et al. 2012). Among the 491 candidate genomic features, 373 features were derived from genomic sequences, 94 features were derived from mRNA expression, and 24 features were derived from molecular interaction. Moreover, the best AUC value was 0.782. In 2015, Lloyd et al. investigated the relationship between phenotype lethality and gene function, copy number, duplication, expression levels and patterns, rate of evolution, cross-species conservation, and network connectivity, and the random forest-based model used in this study achieved an AUC of 0.81, which is significantly better than that obtained by random guessing. In addition, these authors integrated the features they identified to predict essential genes in *A. thaliana* (Lloyd et al. 2015). Those previous researches in three eukaryotes illustrated classifiers can gave satisfactory prediction through combining sequence with other features. For human essential gene identification, we only used the sequence composition and interval association information in the present study and still obtained an AUC of 0.8854. Considering that this result is better than the results obtained in previous studies using integrated features, the gene essentiality of the human genome can be accurately reflected based on only the sequence information.

## Materials and methods

### Benchmark dataset

The human essential genes in cancer cell lines were identified by three different groups (Blomen et al. 2015; Hart et al. 2015; Wang et al. 2015). The results showed high consistency (Fraser 2015), which further confirmed the accuracy and robustness of these data. We extracted the gene essentiality data from the DEG database (http://tubic.tju.edu.cn/deg/), the updated version of which contained human gene essentiality information. These essentiality annotations serve as the basis for constructing our benchmark dataset. The flowchart shown in Figure 2 illustrates the construction of the positive and negative dataset.

**Figure 2.**
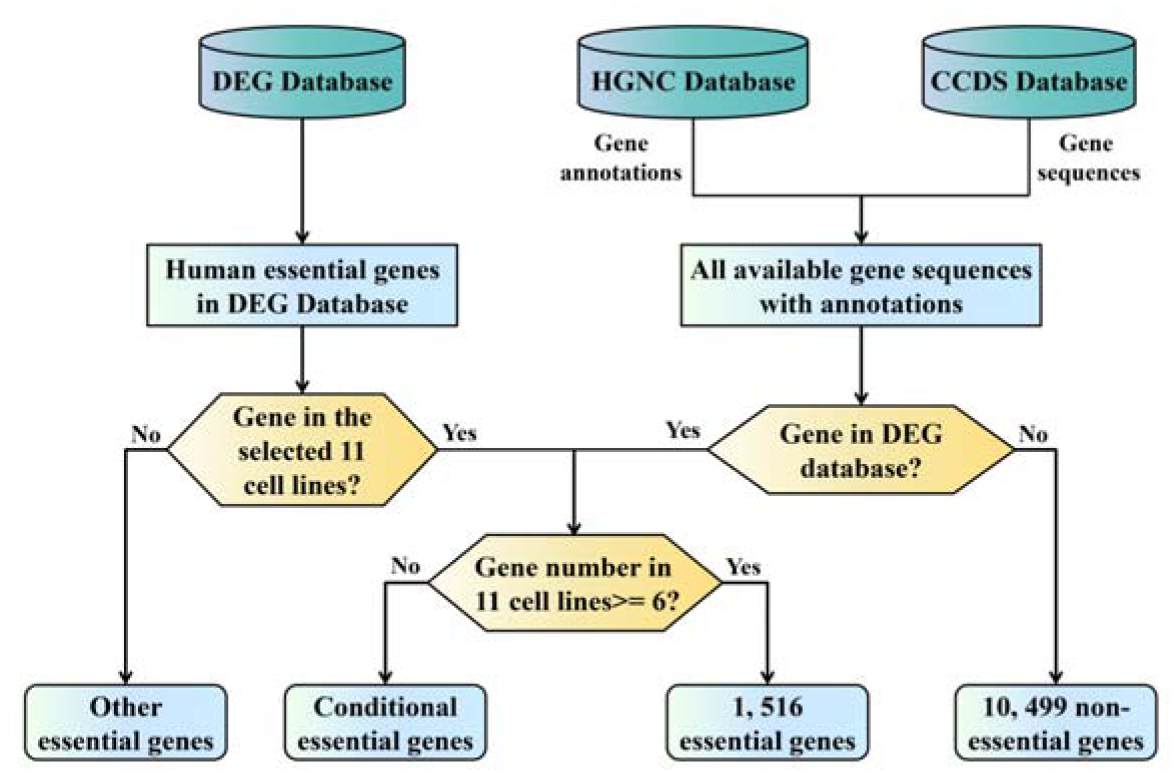
Description of the construction of the human essential and non-essential gene datasets.

In the three studies, 13 datasets were provided. As lines HCT116 and KBM7 are represented by two datasets each, 11 cancer cell lines (KBM7, K562, Raji, Jiyoye, A375, HAP1, DLD1, GBM, HCT116, Hela, rpel) are involved in total. Blomen *et al.* and Wang *et al.*identified the essential genes in the KBM7 cell line. We combined these two datasets into one gene set, KBM7. A total of 2,073 and 386 essential genes were contained in the two datasets for HCT116. The 386 genes in this dataset were markedly different from those in the datasets for the other cell lines, so this dataset was excluded. Ultimately, 11 essential gene sets were obtained, corresponding to a single cell line. Essential genes, by definition, are indispensable for the survival of organisms under optimized growth conditions and are considered the foundation of life (Juhas et al. 2011). Therefore, we only retained genes that were identified as lethal genes in more than half of the investigated cell lines. When a gene appeared as essential in more than six cell lines (11/2≈6), it was selected as one sample in the positive dataset. According to this principle, we obtained a total of 1,518 essential gene annotations. We downloaded all of the protein coding gene sequences from the CCDS database (https://www.ncbi.nlm.nih.gov/CCDS/CcdsBrowse.cgi), and the annotations of protein coding genes were obtained from the HGNC database (https://www.genenames.org/cgi-bin/statistics, March 1, 2016), which contained 19,003 annotation entries. The essential gene sequences were extracted according to the annotations, and genes with no counterpart in the CCDS database were excluded. According to this criterion, we excluded 2 genes and obtained 1,516 essential genes. We used the essential gene annotation in the DEG dataset, and the gene sequences were extracted from the CCDS because the former did not contain the information for non-essential genes. For human essentiality annotations in the DEG database, a number of scattered annotated essential genes aside from those in the 11 cell lines were identified. A total of 28,166 essential gene annotated entries (including conditional essential gene annotated entries) were obtained. Among these annotations, there were many repeated annotation entries; therefore, there were considerably fewer unique entries. To obtain a more reliable negative dataset, i.e., absolutely non-essential genes, we excluded all of the human essential genes annotated in the DEG database (Luo et al. 2014) from the list of the protein coding genes. The remaining genes were regarded as the negative dataset, and their gene sequences were extracted from the CCDS database. Genes with no counterpart in the CCDS database were also excluded. A total of 10,499 non-essential genes were obtained using this method. Ultimately, a total of 12,015 gene entries were obtained in the benchmark dataset: 1,516 essential genes and 10,499 non-essential genes. The protein coding gene annotations are provided in Supplemental Information S5, and information for the benchmark dataset is provided in Supplemental Information S6.

### λ-interval Z-curve

The originally proposed Z-curve variables might reflect the composition of a single nucleotide considering the features derived from phase heterogeneity of a single nucleotide (Zhang and Zhang 1991; Zhang and Chou 1994; Zhang and Zhang 1994; Zhang 1997). Herein, we provided a summary of the Z-curve method used for gene identification (Zhang and Wang 2000). Let us suppose that the frequencies of bases A, C, G and T occurring in an ORF or a gene fragment at positions 1, 4, 7, …, 2, 5, 8, …, and 3, 6, 9, …, are represented by *a*_*1*_, *c*_*1*_, *g*_*1*_, *t*_*1*_; *a*_*2*_, *c*_*2*_, *g*_*2*_, *t*_*2*_; *a*_*3*_, and *c*_*3*_, *g*_*3*_, *t*_*3*_, respectively. Those 12 symbols represent the frequencies of the bases at the 1^st^, 2^nd^ and 3^rd^ codon positions, respectively. According to the symbols defined above, the universal Z-curve mathematical expression is as follows (Zhang and Wang 2000):

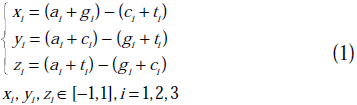

Because composition bias for oligonucleotides in coding DNA sequence (CDS) regions or open reading frames (ORFs) exists, the adjacent w-nucleotides Z-curve method was proposed (Guo et al. 2003; Gao and Zhang 2004). Let us suppose that w represents the length of the adjacent nucleotide sequence. The Z-curve variables for the phase-specific adjacent w-nucleotides can be calculated as follows:

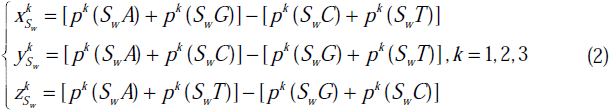

where k equals 1, 2, or 3 to indicate that the first oligonucleotide bases are situated at the 1^st^, 2^nd^ and 3^rd^ codon positions, respectively.

Recent studies demonstrated the existence of long-range associations in chromosomes and showed that these associations are crucial for gene regulation (Fullwood et al. 2009; Ruan 2011). Although the two adjacent nucleotides in the primary structure have no association in some cases, strong associations in terms of tertiary structure might exist. Therefore, we introduced the λ-interval Z-curve to virtually represent the interval range association. The details of this method are described as *p_k_(S_w_X)*, which represents the frequency of oligonucleotides *S*_*w*_*X* in genes or ORFs, where X is one of the four basic bases A, T, G and C. To facilitate this presentation, the length of the oligonucleotide *S*_*w*_ is represented as w. According to the predetermined characters, we generated the universal equation for the λ-interval Z-curve based on Z-curve theory as follows:

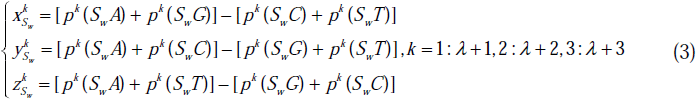

where x, y, and z represent the accumulation of the three base groups classified according to chemical bond properties. Variable k denotes the phase-specific index of the first base in the nucleotide sequence *S*_*w*_, and λ represents the intervals between *S*_*w*_ and *X*. The first base in the oligonucleotide *S*_*w*_ was located at position k. The core part of λ-interval Z-curve forms oligonucleotide *S_w_X.* A schematic diagram of the formation of these oligonucleotides is shown in Figure 3.

**Figure 3.**
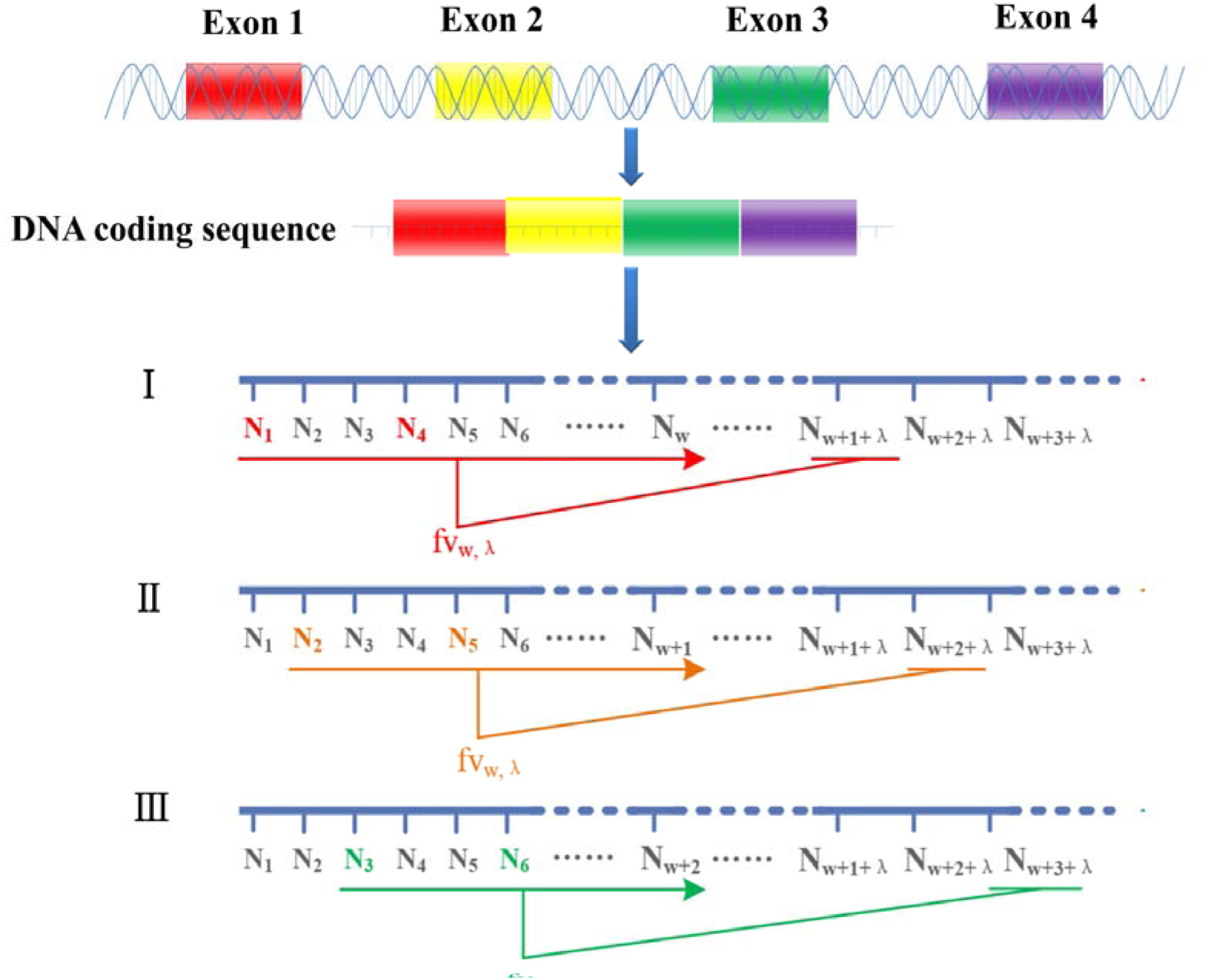
Description of the process of constructing the oligonucleotides using the λ-interval Z-curve theory algorithm. A gene or an ORF has three phases, denoted with different colors, i.e., red denotes the base located in the first phase, yellow denotes the base located in the second phase, and blue denotes the base located in the third phase.

The oligonucleotide window *S*_*w*_ slides along a DNA molecule sequence according to phase, forming oligonucleotide sets with base X, which is λ intervals away from the last base of *S*_*w*_. The periodicity derived from three codons is denoted as *S*_*w*_*X*.

When w is equal to 1 and λ is equal to 0, Equation (3) can be transformed into Equation (1). When w is more than 1 and λ is equal to 0, Equation (3) can be transformed into Equation (2). Thus, the phase-specific single nucleotide Z-curve and phase-specific adjacent w-nucleotide Z-curve are incorporated into the λ-interval Z-curve. Using the λ-interval Z-curve, we can extract more features to characterize DNA sequences. According to Equations (1) and (2), when w is equal to 0, we can obtain 3×3×4^w-1^ variables to characterize DNA sequences. When w and λ are greater than 0, we can obtain 3×3×4^w^ variables. For convenience, we used *fv*_*w,λ*_ to represent the variables with the length of w for oligonucleotides *S*_*w*_, and the highest interval length between oligonucleotides *S*_*w*_ and base *X* is λ. To obtain more information from a DNA sequence, the final variable is described as *FV*_*w,λ*_, where w represents the longest oligonucleotides and λ is the highest interval. This variable can be represented as follows:

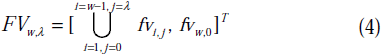

where the symbol ‘U’ represents the union set of *fv*_*w*,λ_, i.e., *FV*_*2,0*_=[*fv*_*1,0*_, *fv*_2,0_]^T^, *FV*_*2,1*_=[*fv*_*1,0*_ ∪ *fv*_*1,1*_, *fv*_*2,0*_]^T^, *FV*_*2,2*_=[*fv*_*1,0*_ **∪** *fv*_*1,1*_ **∪** *fv*_*1,2*_, *fv*_*2,0*_]^T^, …, *FV*_*3,0*_=[*fv*_*1,0*_ **∪** *fv*_*2,0*_, *fv*_*3,0*_], ….*FV*_*2,0*_ and *FV*_*3,0*_ are the combination of adjacent phase-specific w-nucleotide Z-curve variables. We performed this prediction with w ranging from 2 to 4 and λ ranging from 0 to 5. According to the discussion above, we obtained 4,545 variables for *FV*_*4,5*_.

### Support vector machine

SVMs can be classified into linear SVMs and non-linear SVMs according to complexity. SVMs can be used to transform the data into another feature space with more dimensions than the original data and determine the hyper-plane. In this super-dimensional space, the hyper-plane can readily separate the samples. Linear SVMs play a key role in solving ultra-large-scale data, reflecting the effectiveness, rapid speed and splendid generalization of this method in training and prediction. LIBLINEAR, designed by Fan et al. (Fan et al. 2008), is an easy-to-use, freely available software tool to manage large sparse data. The new version of LIBLINEAR (version 2.1-4) supports not only classification, such as L2-loss and L1-loss linear support vector machine, but also regression, such as L2-regularized logistic regression. Given the ultra-high-dimensional feature vectors and large samples contained in the benchmark dataset in the present study, we used the LIBLINEAR software package for prediction. The penalty parameter c was determined using 5-fold cross-validation from 2^−18^ to 2^10^. The new version of LIBLINEAR can be downloaded from https://www.csie.ntu.edu.tw/~cjlin/liblinear/.

### Feature extraction technology

First, the predictive power of a classifier can be influenced by the relevance and noise in the original features. Second, additional time for training and predicting tasks can be increased, reflecting the high-dimensional features. Feature selection (FS) technology is a powerful method for the removal of noise and redundant features from the original features. Hence, the dimension of the features can be reduced. Recursive feature extraction through SVM linear kernels is a powerful FS algorithm (Guyon et al. 2002), but the correlation bias was not considered using this method. Yan and Zhang (Yan and Zhang 2015) proposed an improved method, called SVM−RFE+CBR, which incorporates the CBR. The main concept is that the ranking criterion can be directly derived from the SVM-based model. The feature with the smallest weight is excluded for each run time. The training process was repeated by incorporating CBR until the ranks of all features are obtained. We used SVM-RFE+CBR FS technology to perform feature selection and improve the performance of the classifier.

### Cross-validation and jackknife tests

Cross-validation test technology is the most popular method for evaluating the performance of classifiers. N-fold cross-validation means that the samples are randomly separated into N sub-samples sets. One of the sub-sample sets is used as the testing dataset, and the remaining sub-sample set is used as the training dataset during each run. The process is executed N times until every sub-dataset is utilized as the testing dataset. If N equals the number of samples in benchmark dataset, then the N-fold cross-validation can also be called jackknife or leave-one-out cross-validation. In the present study, we used the 5-fold cross-validation test to determine the best penalty parameter and adopted the jackknife test to further assess the predictive power of the classifier. The area under the ROC curve, the AUC, is often used to measure the performance quality of a binary classifier. An AUC of 0.5 is equivalent to random prediction, whereas an AUC of 1 represents a perfect prediction. There is no bias for evaluating the performance of the unbalanced dataset through AUC. Therefore, we adopted the AUC as a cross-validation criterion in the present study.

## Data access

Supplemental data is available at *Genome Res.* Online:

Supplemental Information S1: The 118 predicted essential genes and their sequences;

Supplemental Information S2: Details for the 17 genes with drug and molecule interactions among the 118 predicted essential genes, distinguished from non-essential genes.

Supplemental Information S3: Sequences of the essential genes in the genome of *M. musculus*;

Supplemental Information S4: The information for the 21 genes and their corresponding homologs.

Supplemental Information S5: Annotation of human protein-coding genes;

Supplemental Information S6: Benchmark dataset used in the present study. Supplemental files can also be downloaded from the CEFG group website: http://cefg.cn/Pheg/supplement_info.zip.

## Acknowledgments

The authors thank Dr. Ke Yan for providing the open source of the SVM−RFE+CBR script. The authors also thank Prof. Chun-Ting Zhang for inspiring discussions and providing invaluable assistance. In addition, the authors also wish to express thanks to the funding for the open access charge: The National Natural Science Foundation of China [31470068]; Sichuan Youth Science and Technology Foundation of China [2014JQ0051]; and Fundamental Research Funds for the Central Universities of China [ZYGX2015Z006 and ZYGX2015J144].

## Author contributions

F. B. Guo conceived, designed and coordinated the present study. C. Dong implemented the main computational analysis. C. Dong, H. L. Hua, S. Liu, H. Luo, H. W. Zhang, Y. T. Jin, and K. Y. Zhang constructed the datasets. C. Dong and H. L. Hua generated the figures. F. B. Guo, C. Dong, and H. L. Hua drafted the manuscript. All authors have read and approved the final manuscript.

